# Nonlinear Singular Value Decomposition Beamforming for Ultrasound Imaging of Gas Vesicles

**DOI:** 10.1101/2024.10.14.618186

**Authors:** Ge Zhang, Mathis Vert, Mohamed Nouhoum, Esteban Rivera, Nabil Haidour, Anatole Jimenez, Thomas Deffieux, Simon Barral, Pascal Hersen, Claire Rabut, Mikhail Shapiro, Mickael Tanter

**Author notes:** Correspondence: Mickael Tanter. These authors have contributed equally to this study (Ge Zhang, Mathis Vert).

## Abstract

Ultrasound imaging holds significant promise for the observation of molecular and cellular phenomena through the utilization of acoustic contrast agents and acoustic reporter genes. Optimizing imaging methodologies for enhanced detection represents an imperative advancement in this field. Most advanced techniques relying on amplitude modulation scheme such as cross amplitude modulation (xAM) and ultrafast amplitude modulation (uAM) combined with Hadamard encoded multiplane wave transmissions have shown efficacy in capturing acoustic signals of gas vesicles (GVs). Nonetheless, uAM sequence requires odd- or even-element transmissions leading to imprecise amplitude modulation emitting scheme, and the complex multiplane wave transmission scheme inherently yields overlong pulse durations. xAM sequence is limited in terms of field of view and imaging depth. To overcome these limitations, we introduce an innovative ultrafast imaging sequence called nonlinear singular value decomposition (SVD) beamforming. Our method demonstrated a contrast imaging sensitivity comparable to the current gold-standard xAM and uAM, while requiring 4.8 times less pulse transmissions. With similar number of transmit pulses, nonlinear SVD beamforming outperforms xAM and uAM in terms of an improvement in signal-to-background ratio of + 4.78 ± 0.35 dB and + 8.29 ± 3.52 dB respectively. Additionally, our method provides a higher flexibility in terms of the selection of acoustic pressure amplitude compared to the other methods. Furthermore, it shows a significant potential for application in the realm of ultrasound localization microscopy (ULM), where it stands poised to facilitate the more precise extraction of nonlinear signatures originating from contrast agents.

## 1. Introduction

Ultrasound plays a pivotal role in biomedical imaging, providing non-invasive, real-time visualization with high spatial and temporal resolution [1]. The recent development of acoustic reporter genes has significantly expanded the capabilities of ultrasound imaging for observing cellular processes and molecular interactions [2]. Acoustic biomolecular contrast agents, commonly known as gas vesicles, have demonstrated substantial potential in a wide range of biomedical applications [3]. These gas vesicles are typically filled with air and encapsulated by a 2-nm-thick protein shell [4], possessing respective diameter and length of approximately 85 nm and 500 nm [5]. Previous studies have revealed that certain types of gas vesicles exhibit a strongly nonlinear response to acoustic pressure (e.g. 0.2 – 0.6 MPa), resulting in nonlinear backscattering of ultrasound waves [6]. This nonlinear behavior enables enhanced detection sensitivity and specificity of acoustic contrast agents, such as gas vesicles, utilizing various nonlinear ultrasound imaging paradigms like amplitude modulation [7].

Both the parabolic amplitude modulation (pAM) [8] and cross-propagating amplitude modulation (xAM) [9] pulse sequences entail line-by-line transmissions of the imaging object, requiring three pulses with relative amplitudes of 1/2, 1/2, and 1 to be pre-calibrated in advance and transmitted during imaging [10]. However, both pAM and xAM pulse sequences are constrained by their imaging depth and framerate, limiting their ability for capturing fast nonlinear events and cancelling tissue motion artifacts, especially within deep tissues. In response to these limitations, Rabut et al. introduced ultrafast amplitude modulation (uAM), an imaging technique amalgamating amplitude modulation, multiplane wave transmission, and selective coherent compounding [11]. This innovation enables the acquisition of nonlinear images after successive transmission of tilted, amplitude-modulated plane waves. It has been demonstrated that the uAM method is capable of achieving ultrafast acquisition of larger and deeper fields of view compared to the other existing techniques for imaging gas vesicles. Nevertheless, this method necessitates odd- and even-element transmissions to generate pulses of relative amplitudes 1/2. Additionally, the imaging scheme involving multiplane wave transmission not only amplifies the complexity of transmission and reception but also strongly elevates the spatial peak temporal average intensity (I_SPTA_) due to prolonged transmit pulses, even at the same mechanical index. Therefore, there is a compelling need to streamline the ultrafast nonlinear imaging scheme to facilitate seamless integration into current ultrasound imaging systems.

To address these challenges, we introduce ultrafast nonlinear singular value decomposition beamforming, a nonlinear imaging paradigm inspired by singular value decomposition beamforming for ultrafast ultrasound imaging [12]. In the last decade, the use of singular value decomposition of ultrasound raw data has been shown to outperform most conventional filtering approaches for the improvement of ultrasound images. First, in linear acoustics, it was shown to discriminate tissue motion from blood flow in ultrafast ultrasound datasets based on spatio-temporal coherence [13] leading to ultrasensitive Doppler imaging. Then, singular value decomposition (SVD) processing of ultrafast compounded plane wave acquisitions was also shown able to improve ultrasound B-mode image quality in the presence of aberrations by estimating the optimal aberration correction law required to produce ultrasound images similar with different plane wave transmit angles [12] or different diverging waves transmissions [14]. Finally, SVD beamforming was applied successfully to retrieve sound speed estimates in ultrasound data and applied successfully in the framework of liver steatosis [15, 16]. SVD beamforming can also be applied to recombine ultrasound images acquired with distinct probes [17, 18]. In the field of nonlinear imaging, higher-order SVD of ultrasonic signals acquired at different pressure levels and different temporal acquisitions was recently proposed to detect moving microbubbles [19]. However, it still relies on the flowing contrast signals. Here, we propose a new nonlinear SVD beamforming approach that does not require any assumption on the motion of contrast agents or gas vesicles. This imaging sequence acquires frames across various acoustic pressure levels. Subsequently, all the frames acquired at the selected pressure levels are concatenated in a Casorati matrix form for SVD processing. Specific modes of the corresponding singular vectors are then selected to reconstruct the final image with high-contrast signals.

The objective of this study is to develop an ultrafast nonlinear SVD beamforming technique that is anticipated to significantly streamline the transmission pulse sequence while achieving better contrast imaging sensitivity and significantly lower acoustic transmission energy compared to the state-of-the-art uAM and xAM imaging schemes. In this study, simulations based on a simple model of backscattered ultrasonic signals in the presence of gas vesicles, speckle noise and background noise were firstly conducted to demonstrate the limitations of the amplitude modulation scheme and the feasibility of utilizing SVD for nonlinear imaging. Subsequently, an *in vitro* gas vesicle phantom was fabricated to validate the pulse sequence and the corresponding post-processing strategies. Signal-to-background ratio (SBR) were employed as image evaluation metric to assess the contrast signal in comparison to the uAM and xAM methods.

## 2. Materials and Methods

### 2.1. Simulation

The phantom was simulated using Matlab (Matworks, USA), as depicted in Figure 1. The final simulated in-phase quadrature (IQ) data, is basically comprised of three matrices describing the spatial distribution of tissue signal, *S*_*Tissue*_, gas vesicle signals, S_GV_, and random noise signals, S_Noise_, respectively. The backscattering amplitudes of gas vesicles, B_GV_, and tissue signals, B_Tissue_, were simulated with respect to the pressure amplitudes, p. Thus, the IQ data can be mathematically expressed as function of spatial and pressure variables:

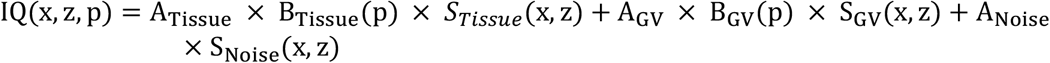

where A_Tissue_, A_GV_, and A_Noise_ are the scalars which represent the amplitude values of tissue, gas vesicle, and noise components, respectively. B_Tissue_ and B_GV_ are the amplitude responses to pressure ramp along the curves which demonstrated the relationship between backscattering amplitude and pressure amplitude as shown in Figure 1(e). It has been noted in prior research that tissue may exhibit a nonlinear signal due to the nonlinear propagation of ultrasound waves within tissues [20].

**Figure 1.**
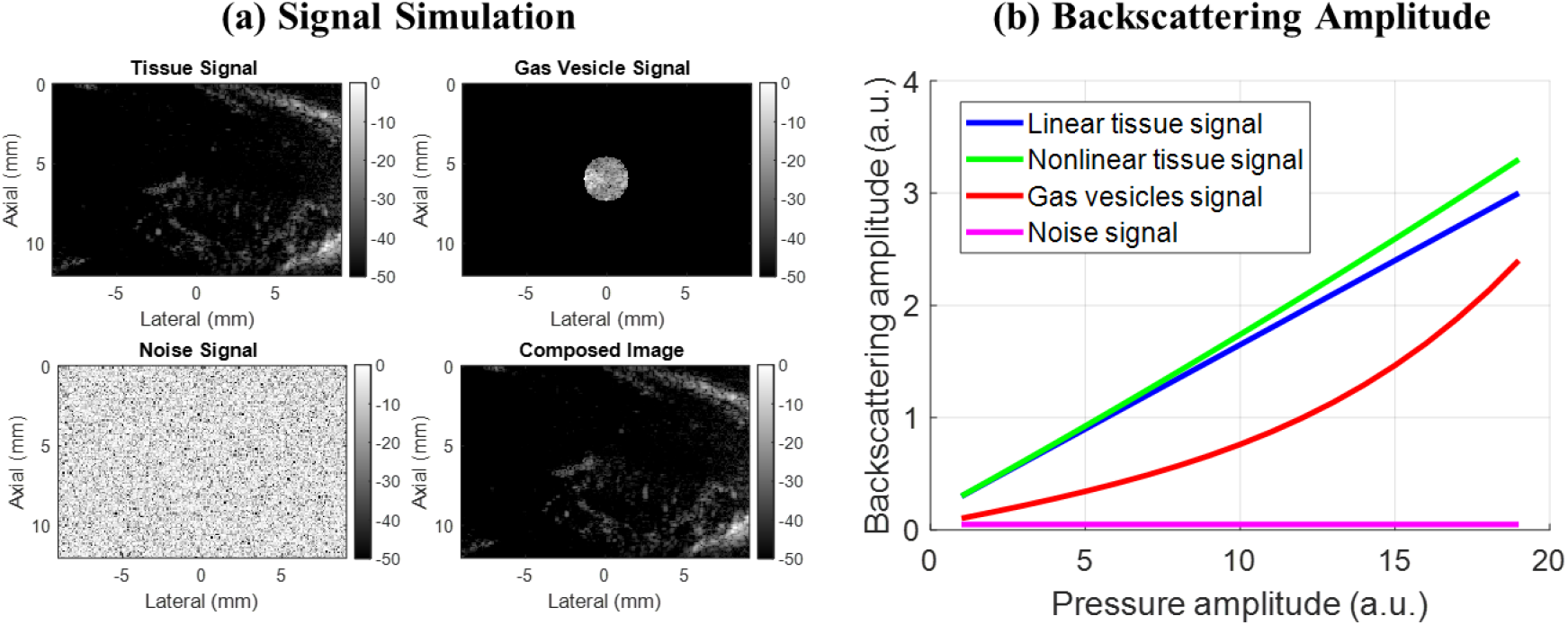
Simulation of ultrasound image composed of tissue, gas vesicle, and noise signals; (b) Backscattering amplitude of tissue signal and gas vesicle signal with respect to pressure amplitude.

The backscattering amplitude of gas vesicle signal, as well as both linear and nonlinear tissue signals, were simulated against various acoustic pressure amplitudes. The response from amplitude modulation at the acoustic pressure amplitude, p can be mathematically expressed as:

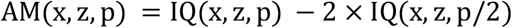

### 2.2. Experimental Setup

Nonlinear GVs were prepared following the methodology outlined in Rabut et al. [11]. In brief, GVs were extracted from buoyant Anabaena flos-aquae cells through hypertonic lysis and subsequently purified by repeated centrifugally assisted flotation and resuspension. The outer GvpC layer of the gas vesicles was then removed through treatment with 6M urea, followed by additional round of centrifugally assisted flotation, dialysis in 1x PBS and resuspension to remove residual GvpC and urea [21].

A tissue-mimicking gas vesicle phantom for imaging was fabricated by casting 1% (w/v) agarose in PBS supplemented with 0.2% (w/v) Al_2_O_3._ For static imaging, a custom 3D-printed mold was employed to create a cylindrical well with a 2 mm diameter. Gas vesicles were incubated at 60 °C for 1 minute, mixed in a 1: 1 ratio with molten agarose, resulting in a final gas vesicle concentration equivalent to 3 OD500nm, and loaded into the phantom. The Al_2_O_3_ concentration was carefully selected to match the scattering echogenicity of the gas vesicle well. This well was precisely centered at a depth of 6 mm. The schematic diagram for *in vitro* setup can be seen in Fig. 2(a).

**Figure 2.**
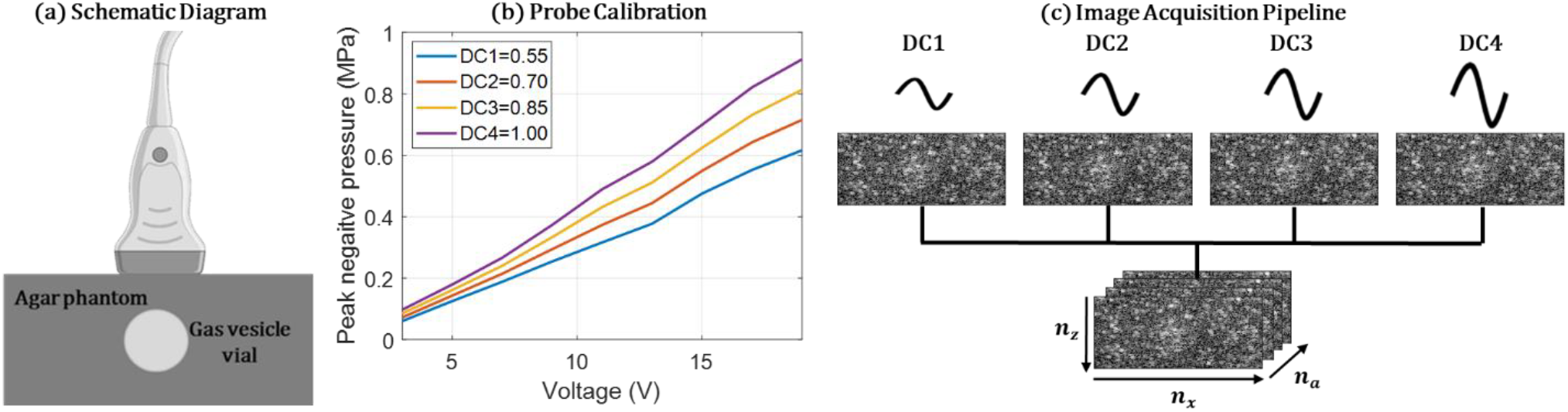
(a) Schematic diagram of the *in vitro* experimental setup; (b) Experimental calibration of the probe used for transmitting pulses with four different duty cycles used for the definition of the transmit signal; (c) Schematic diagram of pulse sequence applied for the image acquisition. Four images acquired with four different duty cycles were used for further image processing.

### 2.3. Data Acquisition

For image acquisition, ultrasound sequences were implemented and executed on a research ultrasound system (Verasonics, USA) driving a linear ultrasound transducer (128 elements, 15.625 MHz central frequency and a 67% bandwidth at -6 dB). All acquisition scripts and processing codes were developed in Matlab (Matworks, USA). The peak-negative-pressure was increased from 0.1 to 0.8 MPa for the transmission pulses based on the calibration result displayed in in Fig. 2(b).

xAM and uAM pulse sequences utilized in this study for comparison with nonlinear SVD beamforming have been visually represented in Figure 3. The number of pulses and transmissions between three different imaging sequences were compared and demonstrated in Table 1.

**Table 1.** Comparison of the number of pulses and transmissions between different pulse sequences. N represents the number of pressure amplitude utilized in nonlinear SVD beamforming.

| Sequence type | Number of waveforms | Number of transmissions | Number of pressure amplitude | Hadamard Encoding | Number of pulses |
| --- | --- | --- | --- | --- | --- |
| SVD (N=4) | 1 | 11 | 4 | 0 | 44 |
| SVD (N=16) | 1 | 11 | 16 | 0 | 176 |
| uAM | 3 | 8 | 1 | 8 | 192 |
| xAM | 3 | 64 | 1 | 0 | 192 |

**Figure 3.**
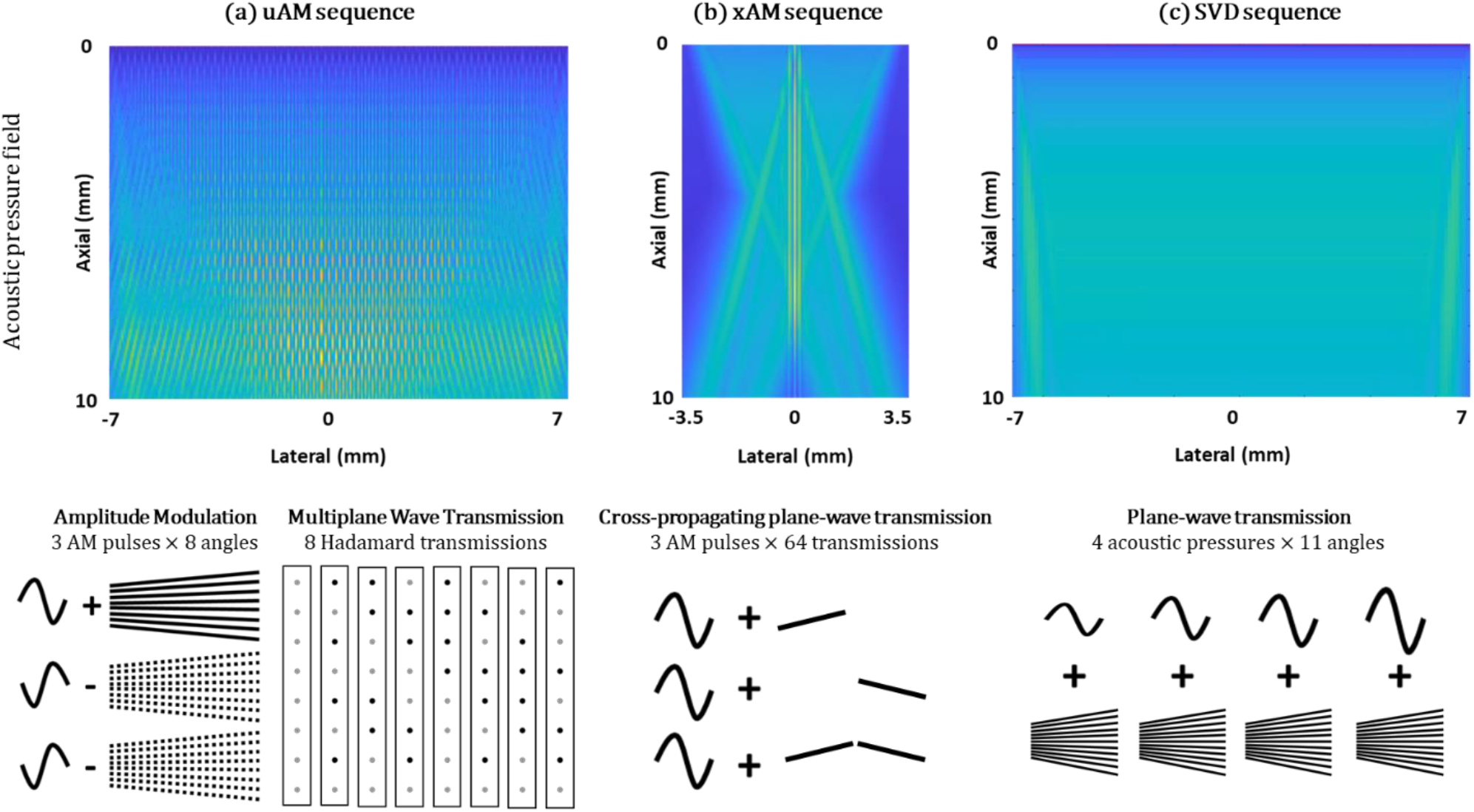
Illustration of (a) uAM pulse sequence, (b) xAM pulse sequence, and (c) nonlinear SVD beamforming pulse sequence, respectively.

In the uAM pulse sequence as shown in Figure 3(a), eight successive tilted plane waves with a transmission frequency of 15.625 MHz were repeated three times with modulated amplitude: two pulses at half amplitude (achieved by silencing the odd and even elements of the transducer, respectively) and one pulse at full amplitude. These sets of modulated pulses were then reiterated eight times. For each repetition, the polarities of the successive plane waves were determined by the column of the Hadamard matrix of order eight.

In the xAM pulse sequence as shown in Figure 3(b), the xAM splits all the 128 elements into two sub-apertures, the first half sub-aperture (element 1-64) and the second sub-aperture (element 65-128). First, the first sub-aperture was used to transmit a tilted plane wave with a transmission frequency of 15.625 MHz at an angle of 19.5° with respect to the array which was optimized in the previous study [9]. Then the second sub-aperture was used to transmit a symmetric plane wave at the same angle with respect to the array. Finally, the two previous two plane waves were transmitted simultaneously.

In the nonlinear SVD beamforming pulse sequence as shown in Figure 3(c), single-cycle plane waves were transmitted at a frequency of 15.625 MHz, with 11 compounded angles for each pulse with an angle range from -10° to +10°. Then the pulses were repeated for 4 different duty cycles as previously calibrated and shown in Figure 2(b).

After the acquisition of ultrasonic raw data using these three imaging sequences, the radio frequency data were offline beamformed using a delay-and-sum beamformer on GPU, utilizing a resolution grid with a spacing of 0.5λ, resulting in the generation of IQ data.

### 2.4. Nonlinear Singular Value Decomposition Beamforming

The processing pipeline of the nonlinear SVD beamforming technique was developed and evaluated in this study. The acquired beamformed image series, *IQ*, in-phase/quadrature (IQ) data can be rearranged by concatenation of IQ image stacks as *IQ*(*x, z, a*), where *IQ*(*x, z, a*) represents the concatenated IQ data acquired at the different acoustic pressure amplitudes. *x* represents the lateral dimension, *z* represents the axial dimension, *a* represents the number of acoustic pressure amplitudes.

The singular value decomposition beamforming was performed to decompose the reshaped data of frames into a weighted, ordered sum of separable spatio-pressure modes expressed by U, S, and V as:

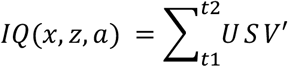

where U and V are the corresponding matrices and S is the weighting matrix. U represent the spatial modes of the acquisition. V represent the pressure modes in this study.

The spatial matrix, U, was used to perform the threshold selection to generate the spatial similarity matrix and square-fitting matrix, respectively, to determine the values of t1 and t2 used in SVD processing as described in the previous study [22]. Briefly, the spatial similarity matrix is obtained from the covariance of modules of the spatial singular vectors, U. Two square-like domains are demonstrated along the diagonal of the spatial similarity matrix, representing the tissue background and the contrast signal subspaces in our study. Then, a square-fitting matrix was converted from the spatial similarity matrix to determine the SVD threshold t1 and t2. Then t1 and t2 were used to filter the weighting matrix, S, to generate a new weighting matrix, S2.

The original S matrix was replaced by the new matrix S2, to generate the SVD-filtered dataset, *g*(*x, z, a*), which filtered out the tissue background signals:

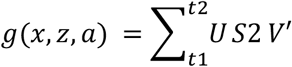

The final image, *g*(*x, z*) was generated by incoherently summing all the images at different acoustic pressure amplitudes as below:

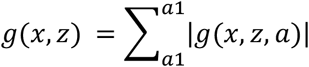

The detailed post-processing pipeline can be seen in Figure 4.

**Figure 4.**
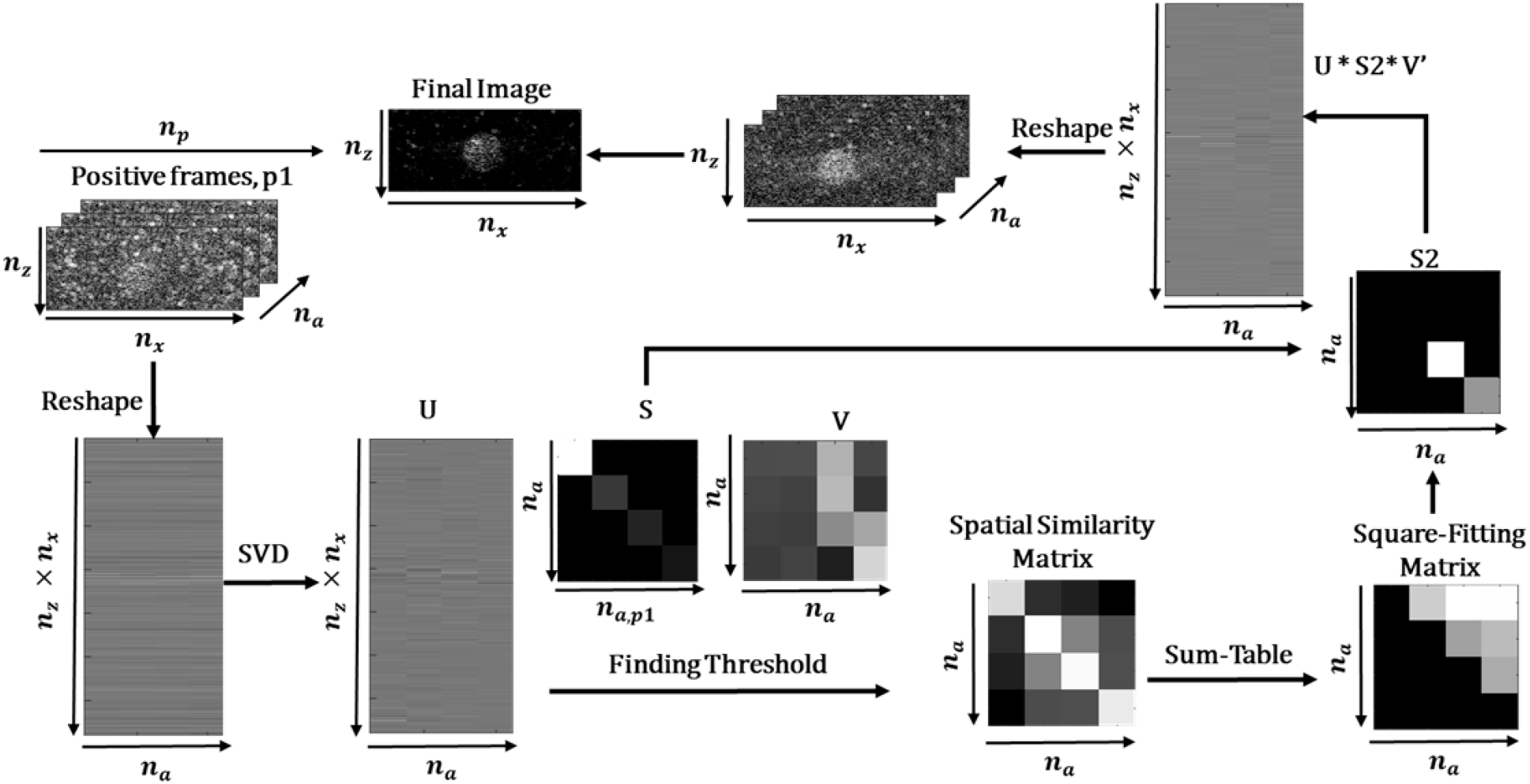
Processing pipeline of the nonlinear singular value decomposition beamforming.

### 2.5. Optimal Selection of Acoustic Pressure

To fairly compare the performance of xAM, uAM, and SVD sequences, each method required to be performed under its optimum imaging pressure. In order to select the corresponding optimum imaging pressure, each sequence was repeated 100 times per second for the data acquisition at a single pressure level. Then the acoustic pressure amplitude was increased progressively ranged from 0.15 to 0.80 MPa to determine when the GVs were destructed for three methods respectively. All the other imaging conditions were kept the same as previously described in the section 2.3. At the end, the optimal acoustic pressure for each sequence (without significant GV destruction) was used for imaging the same GV phantom to evaluate the performance of each sequence.

### 2.6. Image Evaluation

The parameter used for the evaluation of the image quality was signal-to-background ratio (SBR) evaluating the GV contrast as previously described in the literature [10]:

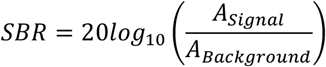

*A*_*Signal*_ is the mean amplitude of signal within the contrast region; *A*_*Background*_ is the mean amplitude of signal within the tissue region.

## 3. Results

### 3.1. Simulation Results

The simulation results presented in Fig.5 show the comparison between the linear and nonlinear tissue signals. It can be seen that the conventional amplitude modulation method cannot completely remove the tissue signal when the tissue exhibits a nonlinear behavior. For the proposed SVD beamforming method, the tissue signals can be significantly removed in both cases of linear and nonlinear tissue behaviors. Fig. 5(c) presents the images obtained by nonlinear SVD beamforming for different ratios between backscattering amplitudes of tissue and GV signals. It can be seen that, the nonlinear SVD beamforming method can better extract the GV signal and suppress tissue signal at a higher ratio between backscattering amplitude of tissue and GV signals.

**Figure 5.**
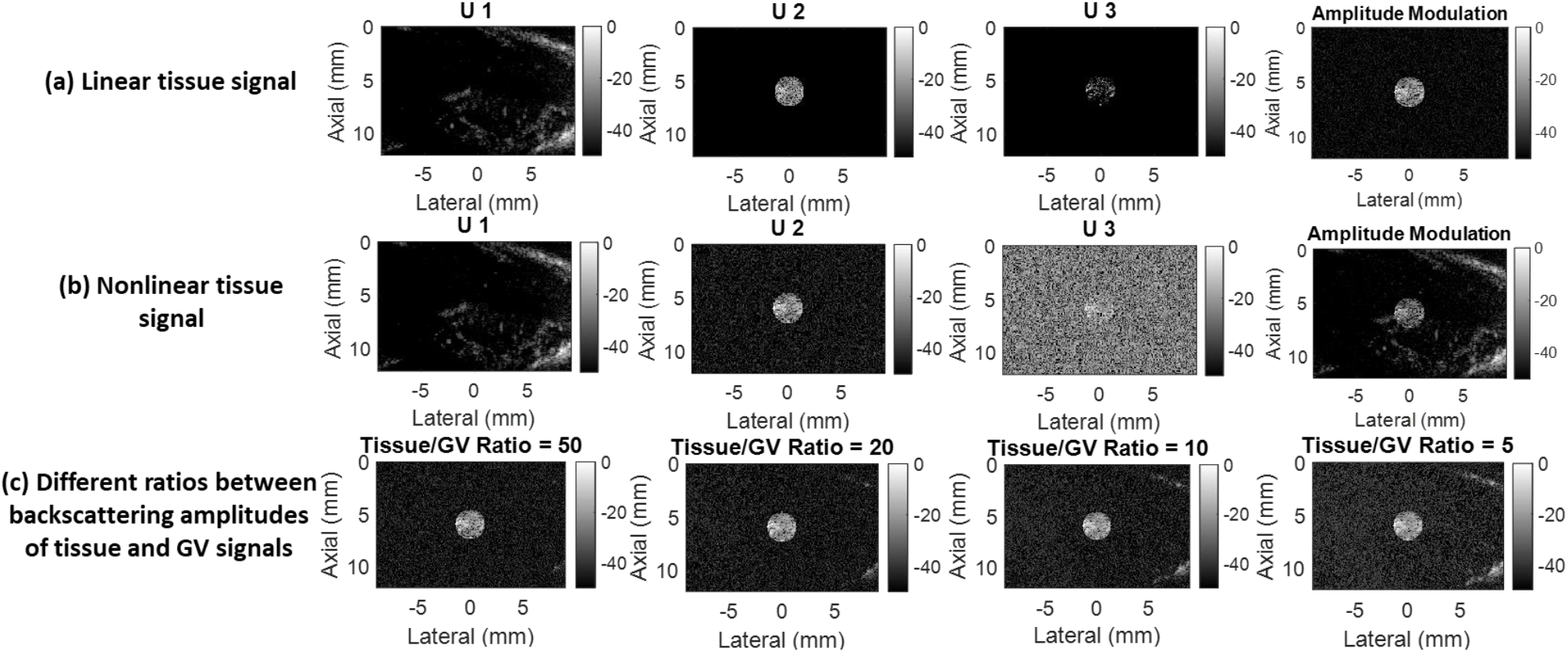
Simulation of nonlinear SVD beamforming on gas vesicle phantom and the first three spatial singular vectors and the corresponding amplitude modulation imaging. (a) when the tissue response is linear to the pressure; (b) when the tissue response becomes nonlinear. (c) The second spatial singular vectors when the ratios between backscattering amplitudes of tissue and GV signals are different.

### 3.2. In Vitro Demonstration of nonlinear SVD beamforming

As can been seen from Figure 6, during the processing of nonlinear SVD beamforming, the input data was decomposed into various modes of spatial matrices, U_i_. Each SVD mode contains different acoustic information which was derived from the original input data. It can be seen that mode 1 represents the tissue signal, i.e. the signal corresponding to highly similar data at different amplitudes. The image obtained in the following modes highlight the nonlinear response and thus the presence of GV inclusions and the nonlinear image artefacts appeared on the modes 2 and 3. In the modes 4 to 6, mainly the GV contrast signals were included. The noise signals are shown in the remaining modes.

**Figure 6.**
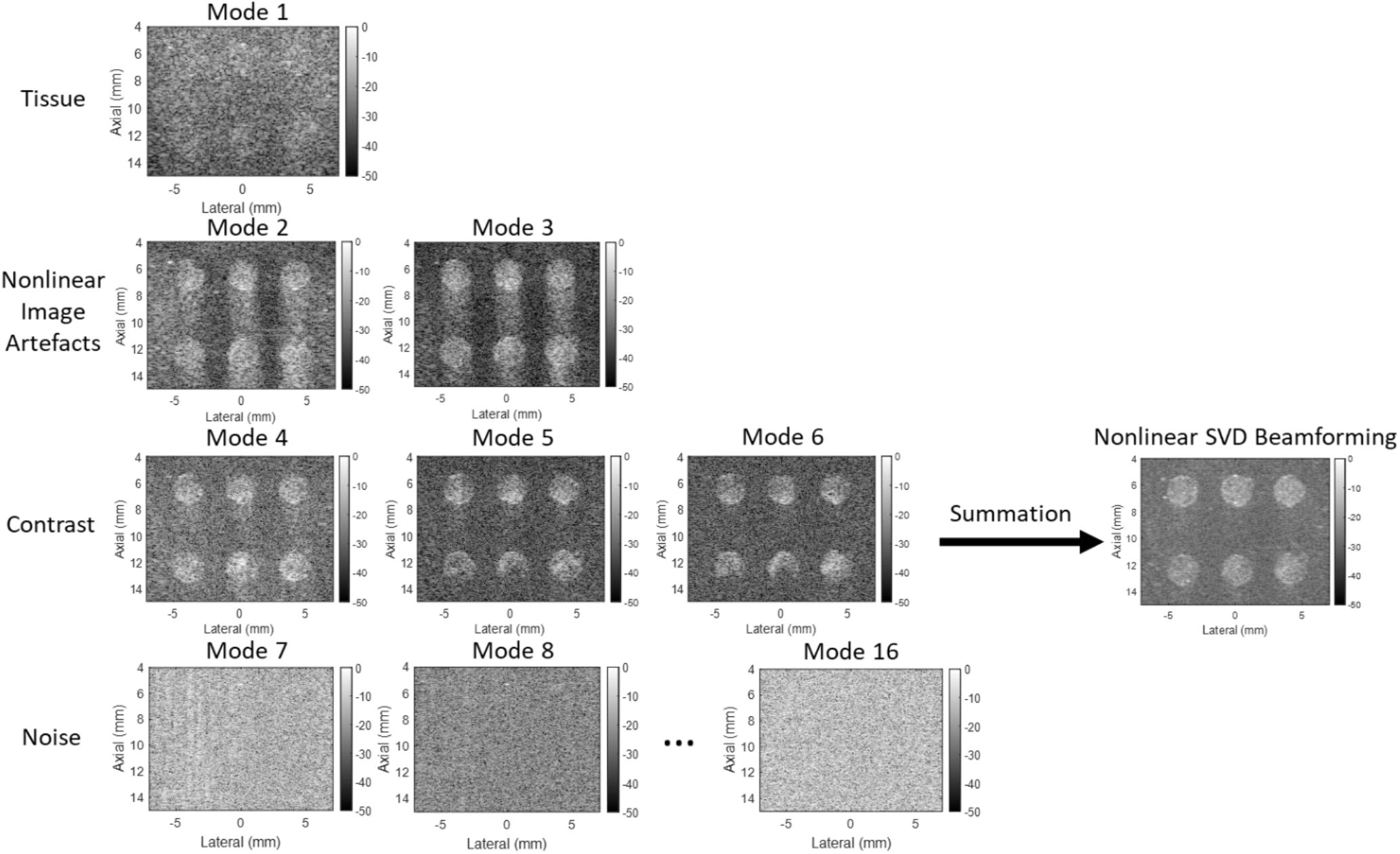
Visualization of the spatial matrix, U for each mode presented in nonlinear singular value decomposition beamforming. Nonlinear SVD beamforming is able to classify the modes into tissue, nonlinear image artefact, contrast, and noise signals. All the images are displayed with a dynamic range of -50 dB.

### 3.3. Dependence on Number of Pressure Level

As can be seen in Figure 7, SBR increased for the simulation results with respect to the number of pressure levels. For the *in vitro* experiments, the SBR level also increases with the number of pressure levels used for the acquisition. Experimental SBR tends to rapidly saturate for pressure levels larger than 4. A N=4 number of pressure levels was used in our further studies as it demonstrates a good contrast imaging sensitivity while it requires a smaller number of transmit pulses.

**Figure 7.**
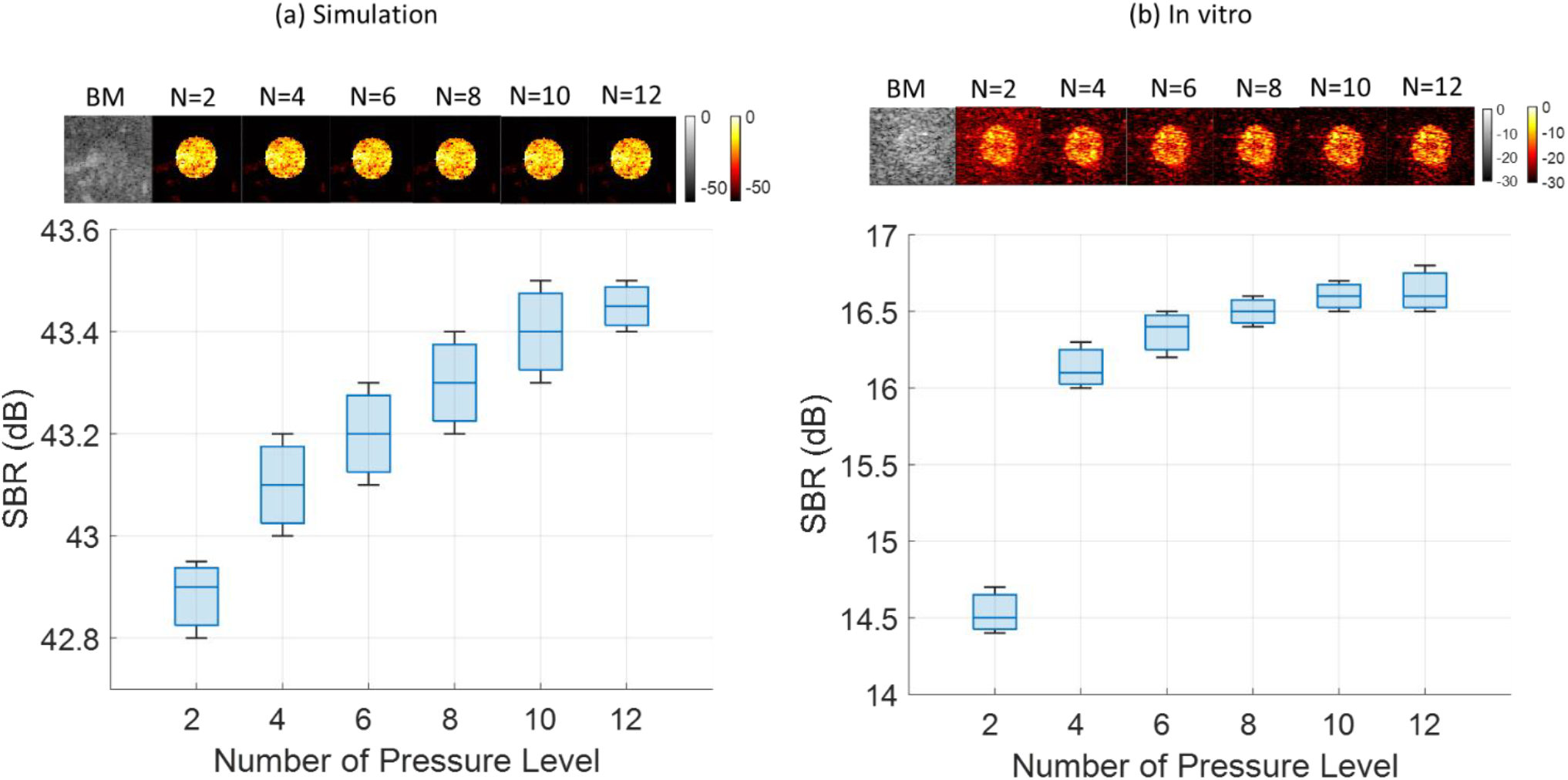
Effect of the number of pressure levels used in the nonlinear SVD beamforming. (a) Simulation images for different numbers of pressure levels; (b) *In vitro* images for different numbers of pressure levels; (c) Image quantification for simulation; (d) Image quantification for *in vitro* experiments.

### 3.4. Optimal Selection of Acoustic Pressure

In order to select the corresponding optimum imaging pressure, each sequence was repeated 100 times per second for the data acquisition at a single pressure level. Then the acoustic pressure amplitude was increased progressively ranged from 0.15 to 0.80 MPa to determine when the GVs were destructed for three methods respectively, which enabled a fair comparison by searching the optimal conditions for each technique. Then the value of SBR was quantified for each sequence at each pressure level along with the visualization of GV contrast.

Figure 8 shows the images the corresponding SBR quantification of the three methods by progressively increasing the acoustic transmit pressures. At each pressure level, the acquisition was repeated for 100 times in 1 second in order to evaluate the potential loss of signal due to GVs progressive disruption. It can be seen that the optimum acoustic pressure required for nonlinear SVD beamforming to obtain the higher SBR is around 0.66 MPa. The SBR value remains high for nonlinear SVD beamforming on a wide range of transmit pressures (0.40-0.94 MPa above 14 dB) compared to uAM (0.65-0.77 MPa above 14 dB) and xAM (0.52-070 MPa above 14 dB). Additionally, the maximal SBR peaks for uAM and xAM appeared to be around 15 dB while our proposed method demonstrated a maximal SBR peak at 16 dB when only four different pressure levels were used.

**Figure 8.**
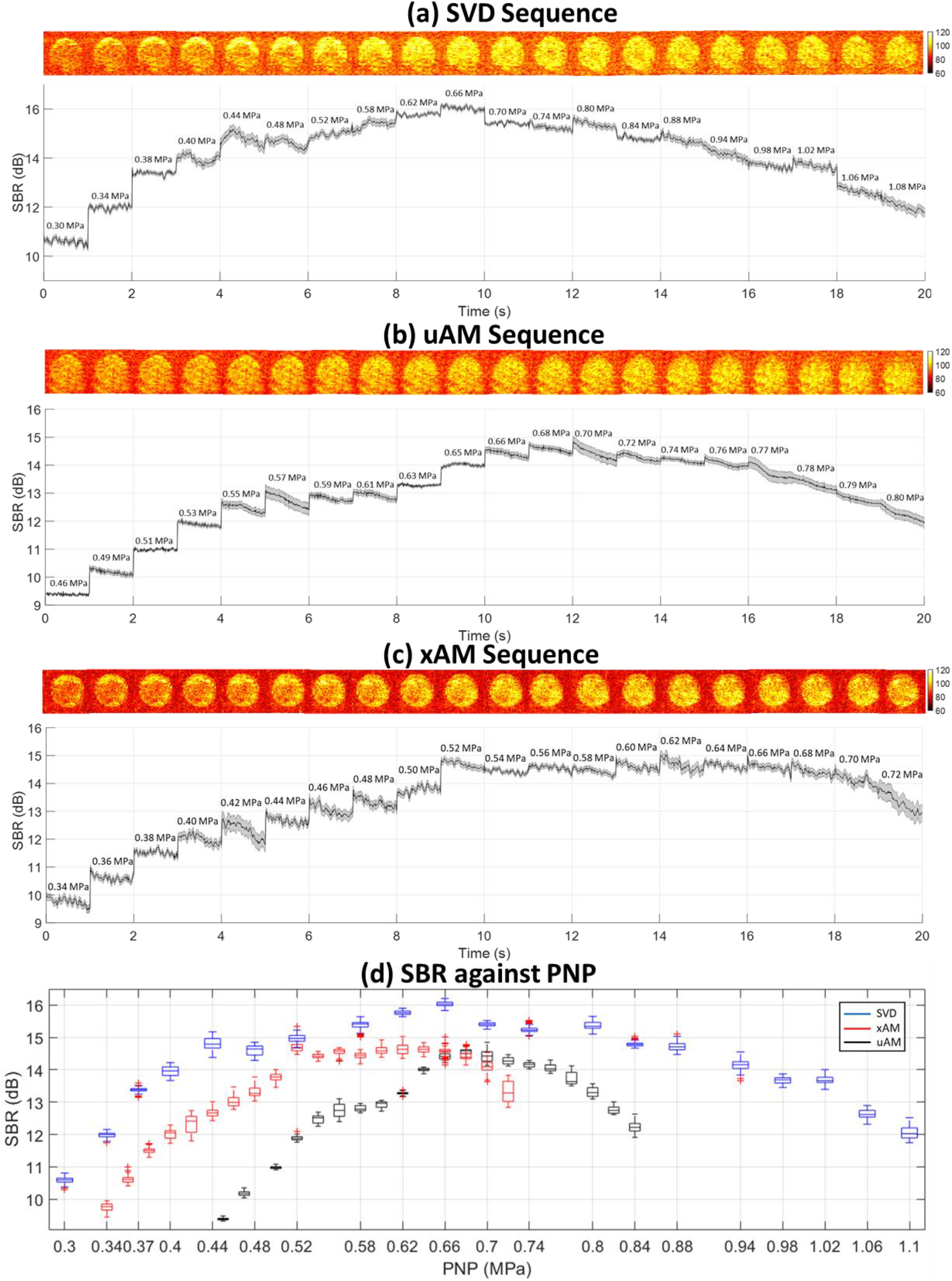
Quantification of signal-to-background ratio (SBR) and visualization of using (a) xAM, (b) uAM, and (c) nonlinear SVD beamforming techniques across the increasing acoustic pressures. The solid line and shaded error bar represent the mean and standard deviation values of SBR correspondingly. All the images are displayed with a dynamic range of -40 dB.(d) SBR across peak-negative-pressure ranged from 0.15 MPa to 0.80 MPa for the comparison of three imaging methods. Red, black, and blue box plots represent for xAM, uAM, and nonlinear SVD beamforming sequences respectively.

### 3.5. Global Comparison between uAM, xAM and SVD sequences at optimal pressure

The results shown in Figure 9 summarize the experimental comparison between xAM, uAM, and nonlinear SVD beamforming methods to extract the GV signals. The optimal acoustic pressures for each imaging methods were found out in Figure 8 and applied for a fair comparison. It can be seen that, uAM image demonstrates a good tissue background suppression capability and a good contrast enhancement of GV signals. However, strong nonlinear imaging artefacts remain present below the GV inclusion. The xAM image demonstrates better axial and lateral resolution and boundary detection of the GV contrast. The nonlinear SVD beamforming method exhibits a higher SBR compared to uAM and xAM methods when the number of pressure level is equal to 4 (Fig 9.c, d, e). For this number of pressure levels, our nonlinear SVD beamforming method required 4.4 times less number of pulses (respectively xx and xx transmits) to achieve such a performance. When a higher number of pressure levels (N=16) was used for nonlinear SVD beamforming in order to reach comparable numbers of transmit pulses, the resulting images were further optimized and provided higher contrast and more tissue background suppression compared to uAM and xAM. The detailed comparison of the number of pulses and transmissions for each method can be seen in Figure 9(f).

**Figure 9.**
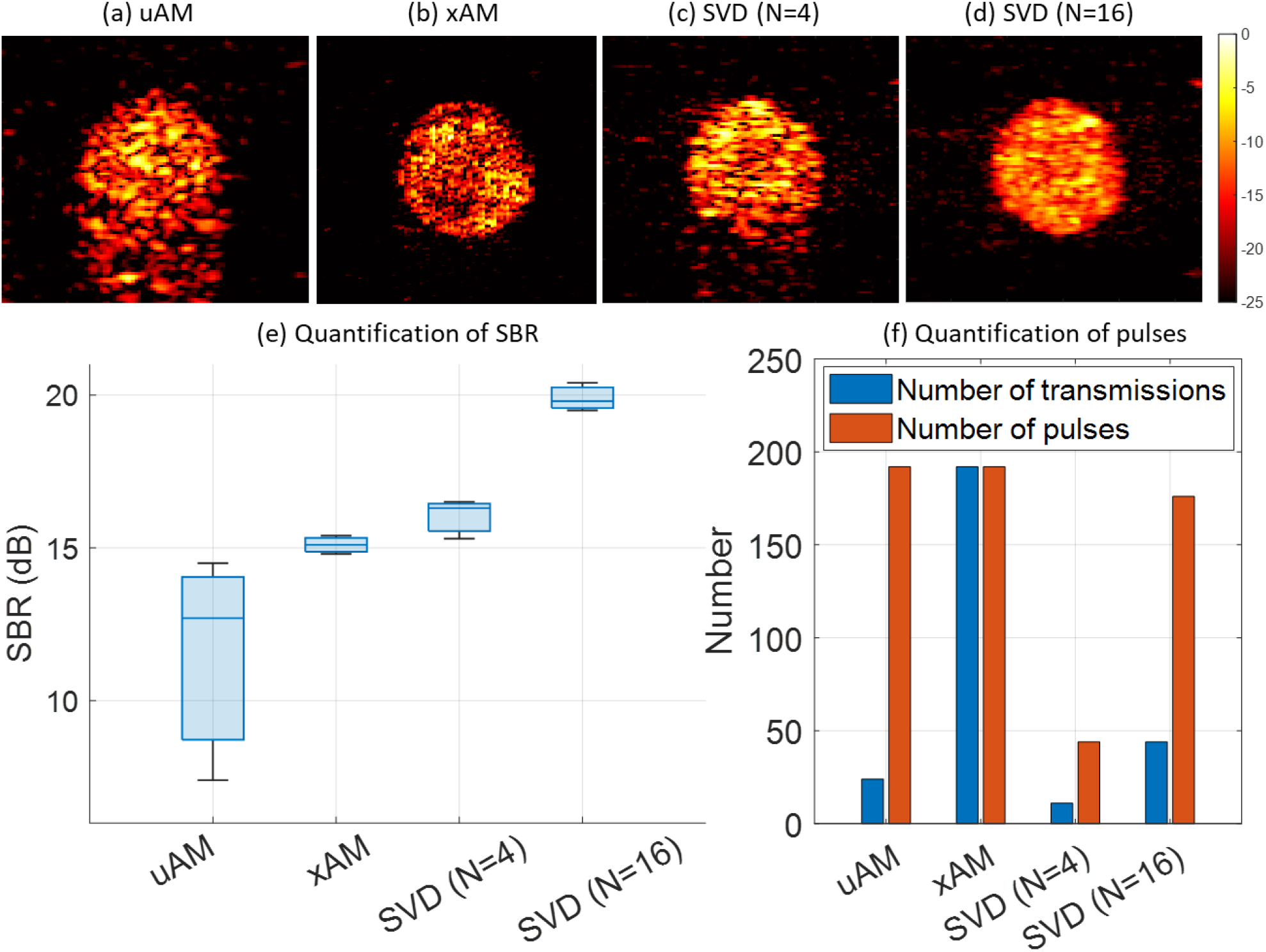
Comparison of *in vitro* GV phantom between uAM, xAM, and our proposed nonlinear SVD beamforming methods. (a) uAM image; (b) xAM image; (c) our proposed nonlinear SVD beamforming method using the number of pressure level of 4; (d) our proposed nonlinear SVD beamforming method using the number of pressure level of 16. All the images were displayed with a dynamic range of -25 dB. (e) Quantification of SBR between different methods. (f) Quantification of the number of transmissions and pulses used for each imaging acquisition.

## 4. Discussion

In this study, we introduce a new nonlinear ultrasound imaging method exploiting the singular value decomposition of ultrafast plane wave compounded data acquired at different pressure amplitude levels. The SVD processing applied to these backscattered ultrasound signals comprising the nonlinear signature of gas vesicles and nonlinear propagation in surrounding tissues extracts independent images, each exhibiting a different nonlinear dependence with respect to the transmit amplitude.. The cancellation of high-order singular value component permits to filter the signals backscattered by tissues and select the nonlinear signals originating from gas vesicles. It is anticipated that our proposed nonlinear SVD beamforming technique will enhance contrast detection and suppress tissue background signals, particularly in scenarios involving strong nonlinear tissue signals and the absence of motion of gas vesicles or microbubbles. The outcomes of both simulations and *in vitro* experiments demonstrate the better contrast detection capabilities of the proposed method while requiring 4.4 times less pulse transmissions or I_spta_ compared to the state-of-art uAM and xAM imaging techniques.

Furthermore, the nonlinear SVD beamforming method helps to remove the nonlinear imaging artefacts which have been previously reported in AM images in the literature [9, 11, 23]. These nonlinear imaging artefacts are inevitable for images based on conventional AM schemes. These artefacts are due to suboptimal ability of AM schemes based on odd/even transmits to cancel linear components of the propagation. It can be seen from Figure 6 that the SVD decomposition process decomposes the spatial modes into four different categories, which are tissue background signals, nonlinear imaging artefacts, GV contrast, and noise. As tissue signals, nonlinear imaging artefacts, and GV contrast signals exhibit different degrees of nonlinearity, SVD processing is able to decompose and rearrange the spatial matrices according to their different degrees of nonlinearity.

In order to provide sufficient sensitivity, the uAM method requires to implement the multiplane wave Hadamard encoding technique for gas vesicle imaging. It involves a notably larger number of pulses and transmissions within the imaging protocol. A prior study demonstrated that, this approach yields a higher spatial peak temporal average intensity (I_SPTA_) owing to the longer transmit pulses [24]. Additionally, it can be seen from Figure 8 that uAM required significantly higher peak-negative-pressure (PNP) to obtain the highest SBR, which may affect its performance *in vivo* when there will be more attenuation of the transmission pressure.

The xAM method exploits the cross propagation of plane waves tilted at a certain angle with respect to the transducer array. Although the xAM image demonstrated a superior image resolution and the boundary detection of GV contrast due to the nature of its focused-shape transmission, its imaging depth, lateral field of view and frame rate are also limited by the intrinsic nature of its transmission scheme. As can be seen from Figure 3(b), the maximum imaging depth is limited at around 8 mm as the angle of sub-aperture plane wave transmission with respect to the transducer array was set as 19.5° as optimized in the previous study [9]. The lateral field of view is limited to the 64 central lines of the ultrasonic B-mode image.

One major advantage of nonlinear SVD beamforming compared to the other conventional AM schemes stands in the fact that the exact transmit pressure amplitude levels do not require to be precisely known and calibrated. Nonlinear SVD beamforming extracts this information without prior knowledge. It is a strong advantage compared to conventional AM transmission schemes that require an exact ½ amplitude transmission. For uAM, the subtraction step faces another additional drawback. The use of odd and even elements transmission in conventional nonlinear ultrasound creates a background noise as the subtraction of odd and event backscattered signals does not reach zero for the linear propagation content.

In this study, a total of four pressure levels were found sufficient to reach high quality images and employed for most data acquisitions as shown in figure 6. Firstly, it demonstrated the highest SBR value among the group. Secondly, it required the fewest number of pulses and transmissions when compared to all the other larger pressure levels and can be achieved at ultrafast frame rates. If sensitivity is favored compared to frame rate, the number of transmit amplitudes can be increased. As shown with 16 transmit amplitudes corresponding to a similar number of transmit pulses compared to xAM and uAM, the imaging sensitivity of nonlinear SVD beamforming is further improved reach even higher image quality (+6 dB compared to AM schemes).

As can be observed in Figure 8 that, the SBR value remains high for nonlinear SVD beamforming on a wide range of transmit pressures (0.40-0.94 MPa above 14 dB) compared to uAM (0.65-0.77 MPa above 14 dB) and xAM (0.52-070 MPa above 14 dB). This demonstrates that nonlinear SVD beamforming is much less sensitive to a non-optimal selection of transmission amplitude. Thus, it provides more flexibility in terms of the selection of transmission amplitude. Additionally, for one selected transmission amplitude, the amplitude will strongly vary *in vivo* due to tissue attenuation in depth. Nonlinear SVD beamforming method will be less sensible to this issue than the other two methods.

The nonlinear SVD beamforming method proposed exploits here the concept of coherence in a new way. The acquisitions and processing permits to discriminate the pixels of the image exhibiting the same dependence with respect to transmit amplitude. Compared to former exploitation of temporal coherence for tissue/blood discrimination [13] and angular coherence for aberration correction [12], the current approach relies of the coherence with respect to transmit amplitude. Our method does not rely on temporal information, therefore it further enables to effectively capture contrast signals of slowly-moving/non-moving gas vesicles either present in tissues or flowing in very small vessels. However, it may also reveal significant promise in the domain of ultrasound localization microscopy [1], poised to enhance the detection of ultrasonic signatures originating from slowly moving contrast agents.

This work has been focused on the development of nonlinear ultrasound imaging of non-disrupting GVs. Further work could adapt and extend this nonlinear SVD beamforming concept to the imaging of GVs disruption. In particular, the exploitation ultrafast data acquired during the fast dissolution curve of GVs just after disruption may be discriminated. Future works will also investigate the in vivo validation of nonlinear SVD beamforming in rodents.

## 5. Conclusions

Our study underscores the potential of ultrafast nonlinear SVD imaging leveraging the capabilities of acoustic reporter genes. The long-lasting pursuit of improved imaging methodologies for *in vivo* detection of contrast agents has driven significant advancements in contrast Ultrasound. This ultrafast nonlinear SVD beamforming technique, utilizing the information from spatial and pressure domains, exhibits compelling performance as demonstrated across simulated and *in vitro* experiments. Nonlinear SVD beamforming technique outperforms other conventional amplitude modulation sequences, even in the complex context of nonlinear tissue responses, while preserving fast frame rate capabilities. Further in vivo studies in rodents should demonstrate the interest of this technique in the context of ultrasound molecular and cellular imaging using acoustic reporter genes.

## Acknowledgement

This work was supported by Inserm research accelerator (Inserm ART) in Biomedical Ultrasound and by the French national research agency (ANR) under ANR-21-CE19-0050 program (Project SonoGT).

## Notes

### Competing Interest Statement

The authors have declared no competing interest.

